# Body size and light environment modulate flight speed and saccadic behavior in free flying *Drosophila melanogaster*

**DOI:** 10.1101/2024.07.08.602594

**Authors:** Elina Barredo, Haoming Yang, John Paul Currea, Yash Sondhi, Ravindra Palavalli-Nettimi, Simon Sponberg, Vahid Tarokh, Jamie Theobald

## Abstract

For flying insects, visual control relies on acquiring adequate light, but many circumstances limit this, such as dim environments, high image speeds, or eyes of modest light gathering power. To determine these effects on vinegar flies, we limited light by either placing them in dim conditions, or generating individuals with developmentally smaller eyes, then examined activity levels and three-dimensional flight paths. When simulating dawn and dusk light periods, walking flies increase activity, reflecting their crepuscular nature, and this is stronger for flies with larger eyes. When light switches abruptly, similar to many lab settings, activity associated with crepuscular periods diminishes, as does activity associated with greater facet size. During free flight, we find flight speed decreases similarly in both dim light and small eye conditions, but excess light induces smaller individuals to restore their flight speed. Through a machine learning approach, we confirmed that two features, translational speed and saccade distance, are sufficient to classify treatment groups by light niche, size, and age. Together, these imply that flight changes in smaller individuals result from visual deficits, rather than other elements of body structure.

## INTRODUCTION

Insects have evolved visual systems that allow them to achieve remarkable flight performance in complex environments. Flies are especially aerobatic (Taylor, 2001), and rely on vision during flight to find food sources, escape predators, avoid obstacles, and stay on a preferred course (Currier et al., 2023; Horridge, 1977; Land & Nilsson, 2012), although these can be highly coordinated with other senses (Huston & Krapp, 2009; Keesey et al., 2019). But limiting light compromises visual behaviors, as insects experience reduced spatial resolution, temporal resolution, and color discrimination (Land, 1997; Sondhi et al., 2021; Theobald et al., 2007; Warrant, 2017). The relationship between light gathering capacity and light availability therefore explains many adaptations in animals with compound eyes, such as nocturnal species with larger facets, which capture more light and compensate for dim conditions (Horridge, 1977; Kapustjanskij et al., 2007; Land & Nilsson, 2012; Snyder et al., 1977).

Holometabolous insects have distinct larval and pupal stages before adulthood, and larval nutrition significantly influences the ultimate size and characteristics of the adult (Currea et al., 2018; Shingleton et al., 2009; Stern & Emlen, 1999). Laboratory-reared insects generally acquire abundant nutrients, generating larger adults than typically found in the wild, where food can be scarce. Since compound eye size is proportional to body size, smaller individuals may have compromised visual acuity and optical sensitivity (Currea et al., 2018; Kapustjanskij et al., 2007). Interestingly, small *Drosophila melanogaster* compensate for their reduced optics in the frontal field by sacrificing temporal resolution to improve contrast sensitivity (Currea et al., 2018), but we do not know how these limitations affect daily activity. For small flies competing with large conspecifics, keeping temporal frequencies within a visible range may require that they (1) restrict activity to brighter locations or times of day to capture more light; (2) or fly more slowly or further from objects to limit optic flow speed.

To test the first prediction, we measured fly activity in a locomotion activity monitor (LAM) under both a daily binary light cycle, that switched intensity abruptly without dim periods, and a gradual light cycle, that simulated the ramped changes in brightness during sunrise and sunset. We then measured mean ommatidial diameters of the flies and compared the overall activity between these conditions. Since adult age can affect behavior and flight ability (Lane et al., 2014; Pithan et al., 2022), we further tracked fly age and its relation to activity. To test the second prediction, we captured flight trajectories (Keller et al., 2020) over multiple days in a restricted light environment, then compared flight characteristics under different illuminations (Fig. 1A, F). We further compared trajectories of flies with smaller eye sizes, which limit light even in bright conditions. Finally, since age, light environment, and size can have non-linear and complex effects on flight performance, we chose a data-driven method to classify how features of flight relate to these conditions. We trained a neural network that could classify fly groups, such as size and light environment, on the basis of flight features.

**Fig. 1.**
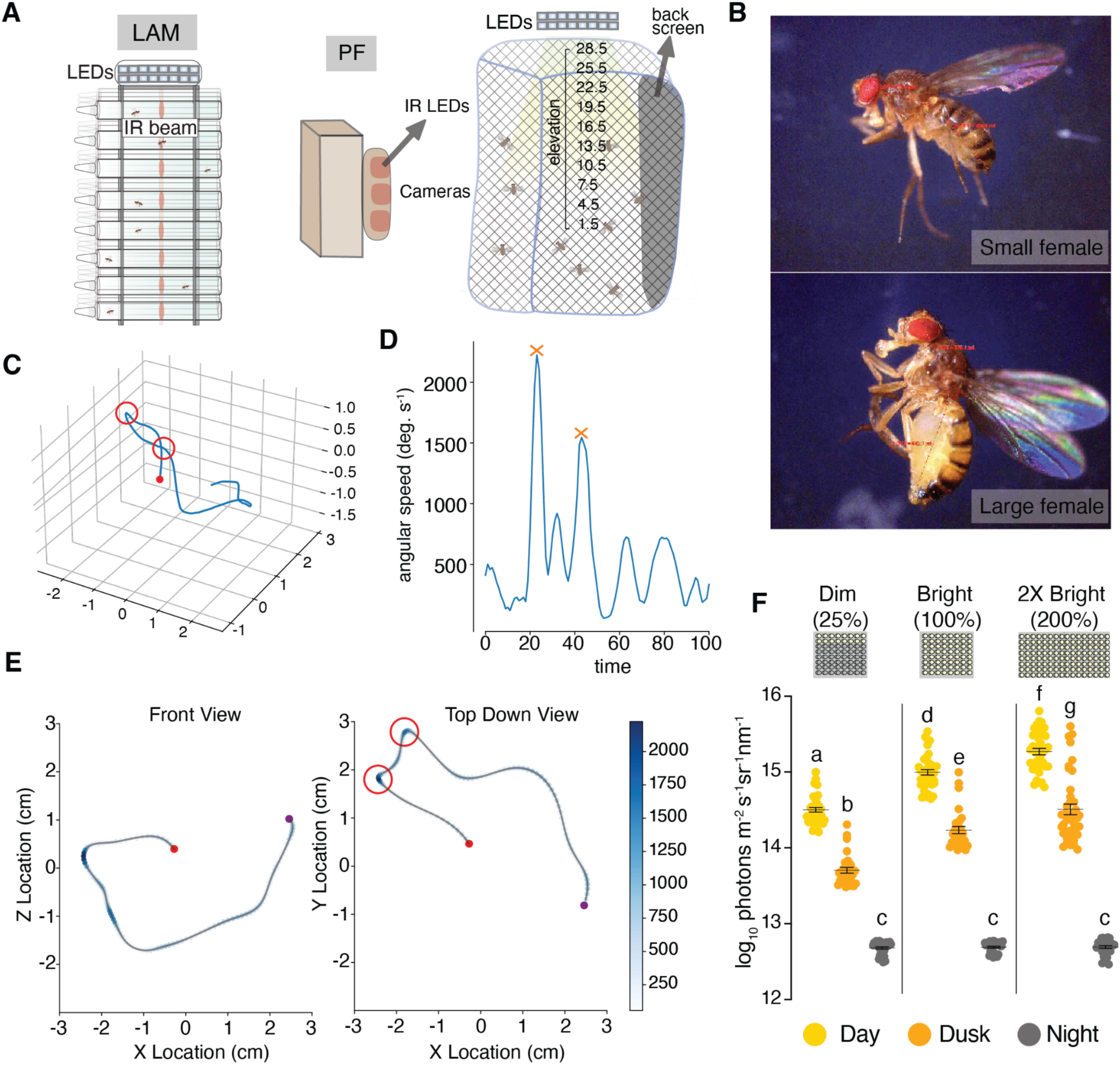
Experimental set-up and sample trajectory. (A) The Locomotor Activity Monitor LAM (left, Trikinetics, Inc) and the Photonic Fence PF (right), a high speed 3D tracking device in front of an insect cage. We varied light environments using a custom controlled LED panel. (B) Body size comparison between a food-restricted (top) and a normally fed (bottom) female fruit fly under a compound microscope. (C) Sample trajectory with two saccadic turns (open red circles). (D) Peaks in angular speed (stars) used to identify saccades in the sample trajectory. (E) Front view and top down view of the representative flight trajectory. Darker blue shaded dots represent faster angular speed in degrees per second. (F) Light level measurements for each lighting condition as estimated using the environmental light field toolbox (ELF, (Nilsson & Smolka, 2021). Lower case letters denote statistical differences (alpha < 0.05, unpaired t tests) between mean light readings for each light environment.

## MATERIALS AND METHODS

### Animal subjects

*Drosophila melanogaster* came from a laboratory colony developed on standard food medium under a 12L:12D light cycle at 21°C. For LAM experiments, we washed third instar larvae through a strainer to remove them from food and generate a broad distribution of body sizes, and therefore ommatidial diameters. We then randomly assigned pupae to one of two LAMs in portable incubators with different lighting schedules, 32 pupae into each of the two conditions. We transferred pupae to tubes containing fresh media 1-2 days before eclosion, and into the LAMs. After recording for 7 days post eclosion, we refrigerated flies overnight, then captured micrograph depth stacks using a digital recording microscope (Zeiss Axio Scope.A1), and obtained ommatidial diameters by applying an automated tool for measuring compound eye parameters. Flies that showed no mean activity on the 6th day were omitted from later analysis, resulting in 23 flies in the abrupt light condition and 21 in the gradual light condition. For all free-flight experiments, we randomly selected 12-15 flies, 4 to 10 days old, of mixed sexes, and transferred them to an experimental cage. For size experiments, we randomly selected the larvae to move to a vial of fresh food or to one containing only a moist cotton roll (see Currea et al., 2018). At adulthood, the foodless flies returned to vials of food media. We confirmed the size differences by measuring body length on ImageJ (Fig. 1B). To test age effects (see Supplementary material), we monitored pupae until adults started to emerge, then selected flies between 12 and 36 hours after emergence. We placed these in a flight enclosure and monitored them for 5 to 7 days.

### Locomotor Activity Monitors (LAMs)

We assessed general locomotor activity using two 32-channel LAMs; LAM 25; 25 mm diameter, 125 mm long; Trikinetics Inc., Waltham, MA, USA) under different light schedules: a conventional abrupt schedule of 12L:12D (on at 7:00 and off at 19:00) and a gradual schedule with light increasing or decreasing linearly over a 2 hour period (ramp on from 6:00 to 8:00 and ramp off from 18:00 to 20:00, represented by the grayscale images at the top of Fig. 2A, B). The full brightness within each incubator ranged from 50 to 2000 lux depending on the position and orientation of our sensor (TSL2591, Adafruit). Both light sequences were generated by a custom system involving a Raspberry Pi, an LED strip, and a custom Python program. The gradual light schedule was designed to keep the same mean and max luminance and circadian period as the abrupt schedule while simulating the gradual change in brightness during sunrise and sunset, based on the shape of data from (Theobald et al., 2007). The LAM quantifies locomotor activity as the number of times a fly interrupts an array of infrared beams, stored as counts per minute. For producing the activity heatmaps in Fig. 2A-D, we smoothed individual fly activity traces using a 60-minute rolling mean and then normalizing to the maximum value of all smoothed fly traces, 1.69 events per minute. The un-smoothed, normalized values were binned for calculating the means in Fig. 2E-G and their corresponding statistics. We defined early activity as occurring between 6:00 and 10:00 and late as between 18:00 and 21:00 (gray spans highlighted in Fig. 2C, D) and used the mean over each entire day for modeling general activity (plotted in Fig. 2F, G).

**Fig. 2:**
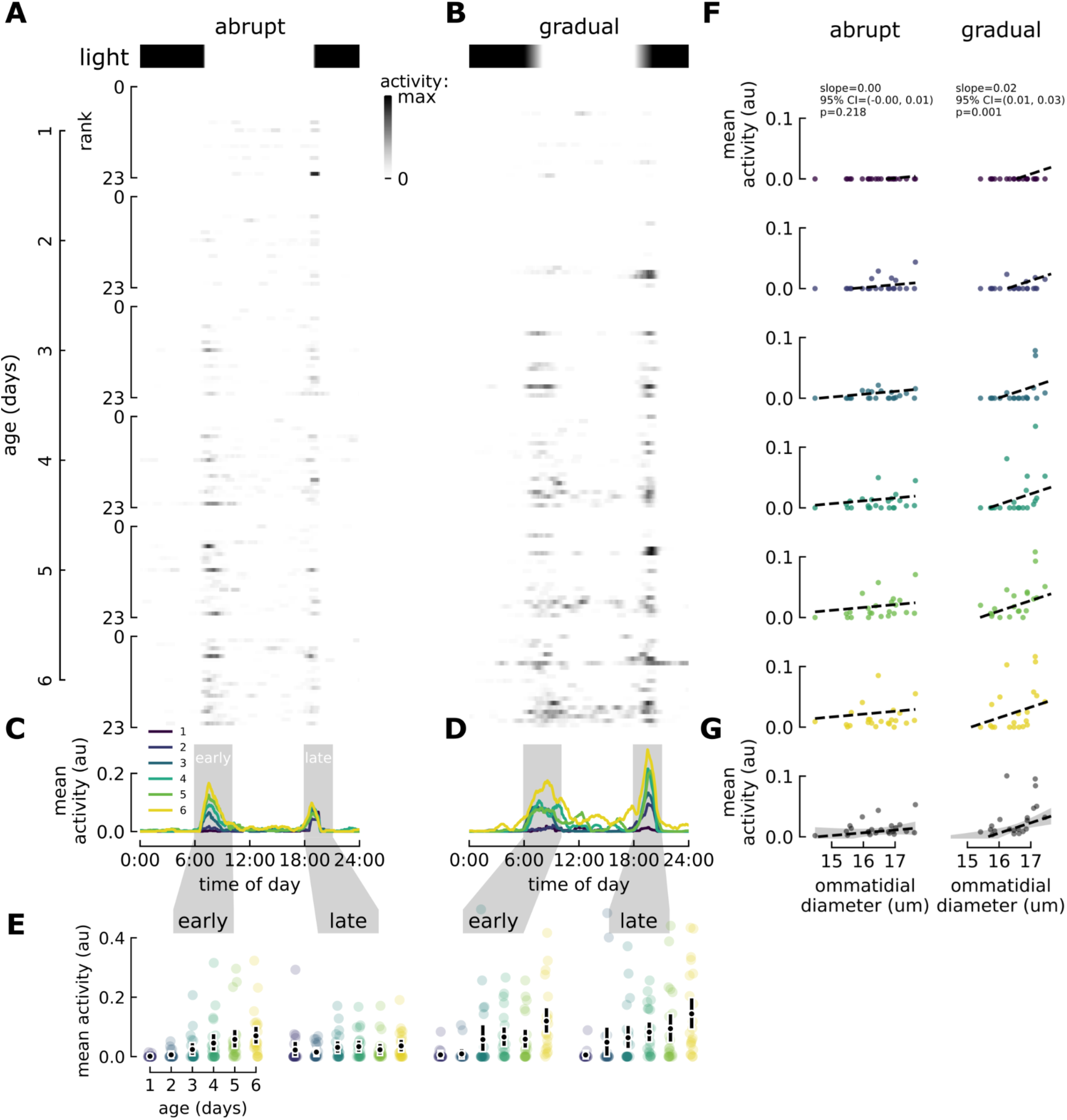
Fruit fly crepuscular activity depends on age, light schedule (abrupt vs. gradual), and ommatidial diameter. (A) Flies reared on an abrupt light schedule (with brightness represented by the grayscale bands at the top). Each row of pixels represents an individual fly’s mean activity smoothed using a 1hr rolling mean and normalized to the maximum smoothed activity of all flies. The rows have been sorted by increasing ommatidial diameter with smallest diameter at the top (rank=1) and largest at the bottom (rank=23). Note that these share the same x-axis as in C. (B) Flies reared on a gradual schedule, with light increasing from 6:00 to 8:00 and decreasing from 18:00 to 20:00, keeping the same mean luminance as in (A) and plotted in the same way. Mean activity in the abrupt light schedule (C) and the gradual light schedule (D) for each day were normalized to the maximum mean activity (1.69 events per minute) and therefore expressed in arbitrary units (au). Spans labeled "early" and "late" highlight crepuscular activity and were used to calculate the means plotted in (E). Error bars represent the bootstrapped 84% C.I. of the mean accounting for within-subject covariances, such that non-overlapping error bars signify statistical differences with an alpha of .05 (Goldstein & Healy, 1995). (F) Mean activity for each day (rows) and light schedule (columns). A GLMM was used to predict means plotted as dashed lines. (G) Overall mean activity by ommatidial diameter per light condition. The dashed line and error bands indicate the mean and 95% C.I. of the mean produced by an OLS model.

### Ommatidial measurements

7 days after eclosion, we transferred surviving flies in both LAMs to individual Eppendorf tubes labeled with their row and column in the LAM, and stored them in a refrigerator overnight. We attached them to thin tungsten rods at the scutum of the thorax for easier manipulation, angled to center the eye on the microscope objective, and then captured depth stacks as in Currea et al. (2018, 2022). We ran these through the ommatidia detecting algorithm (ODA), an open-source Python package for automatically detecting the centers of ommatidia (Currea et al., 2023). The specific code and datasets used to generate the plots and statistics in Figure 1 and Supplemental Figure 1 can be located in a figshare repository (https://doi.org/10.6084/m9.figshare.26122231.v2; Barredo et al., 2024) and the latest version of the ODA can be found here: https://github.com/jpcurrea/ODA.

### Statistical Models of Locomotor Activity

To test the effects of light condition and age on mean daily activity, we used two regression models and bootstrapping. For comparing mean daily activity during the early and late time spans, we bootstrapped to generate an empirical sampling distribution of the mean for each day, sampling all of the measurements per fly in batches 10,000 times with replacement to maintain within-subject covariances. From these we obtained the mean and 84% C.I. of the mean, permitting visual mean comparisons based on the overlap of error bars (Fig. 1E) (Goldstein & Healy, 1995). Then, we took a systematic approach to find a good fitting linear regression model (ordinary least squares, OLS) of mean overall activity as the sum of ommatidial diameter, lighting condition (abrupt vs. gradual), their interaction and all combinations thereof.

Because the lighting condition was categorical, we dummy-coded the relevant regression models. Dummy coding allowed us to include categorical variables in multiple regressions alongside continuous variables (Wolf & Cartwright, 1974) and to therefore test categorical hypotheses about the effect of the lighting condition even after accounting for other variables. In this coding procedure, two different coefficients represented whether the lighting was abrupt or gradual using the abrupt condition as the reference group. For instance, the constant coefficient in model #2 represents the mean daily activity of the abrupt lighting condition and the gradual coefficient represents the difference between the gradual and abrupt lighting conditions. If the gradual coefficient is significant and positive, this means that there was a significant increase in mean daily activity as a result of gradual lighting. For all of these models, ommatidial diameter was mean centered, such that the resulting constant coefficient would serve as a measure of mean activity for the mean ommatidial diameter. Goodness of fit measurements and the resultant coefficients and their p-values are in Supplementary Table 1. We repeated the procedure for early and late activity (for instance, see Fig. S1), but the results were the same qualitatively. Once we arrived at a best-fitting model, we modeled daily mean activity as the sum of condition, the interaction of condition and ommatidial diameter (just like the best-fitting OLS model), and age using a generalized linear mixed model to account for within-subject covariances. The results of this model for general activity are included in Supplementary Table 1 and the predicted means are plotted as dashed lines in Fig. 2F.

### Flight trajectories

We recorded three-dimensional flight trajectories with a Photonic Fence (PF) monitoring device (Mullen et al., 2016). PF uses two infrared cameras and infrared (IR) lights that shine onto a retroreflective screen. As insects in front of this screen occlude the light, the device tracks their position at 100 Hz (Fig. 1A). During trials, we kept flies in a mesh cage (61 x 61 x 91 cm), in a dark room at approximately 22 °C and 40% humidity. Lights above the cage dimmed to simulate dawn from 5:00 to 7:00, and dusk from 17:00 to 19:00. When comparing flies of different size ranges, we placed two insect cages side by side and ran the experiments in parallel. We could then use lateral position to separate cages, but could not test bright and dim light treatments in parallel. We ran at least two biological replicates for each light treatment and included all the trajectories in further analysis. In all experiments, the insect cage contained a flask with fresh food media.

### Environmental light field toolbox (ELF) analysis

To estimate the light levels in the insect cage, we used the Environmental Light Field method as described in (Nilsson & Smolka, 2021). We used a Nikon D850 camera with a Sigma 8 mm/F3.5 lens. We took a dark calibration with a 1/400 s exposure time, 1250 ISO and 3.5 f-stop. For each light condition, we took images at every 5–15-minute intervals from 14:00 to 20:00, obtaining light values before, during and after dusk. We ran the batch mode in the ELF software using the same dark calibration for each image. By default, the ELF method gives the red, green, blue (RGB) and white light intensity in lit (log photons m^-2^ s^-1^ sr^-1^, nm^-1^) for elevations from -90 degrees to +90 degrees, from which we averaged intensity values from 1.5 to 28.5 degrees (see Fig. 1A). We then calculated the mean lit values at three time points: day (16:00), mid-dusk (16:00), and night (20:00), (Fig. 1F).

### Data Preprocessing and statistical analysis

To extract relevant trajectories, we excluded extreme values of speed and distance: any outside the 0- 95%-quantile range, which usually represent localization errors; and those lasting less than one second or with mean speed less than 0.1 m/s, which likely represent walking. We applied the preprocessing routine to all experiments (Supplementary Table 2), then ran two-sided z tests using the statsmodel Python package to compare the means of relevant datasets and determined statistical significance with α=0.05.

### Saccade Identification

Flight in flies includes straight bouts, and at least two distinct types of turns, smooth and saccadic (Land, 1997; Mongeau & Frye, 2017). Body saccades are rapid changes in body direction during navigation that allow insects to reduce retinal distortions that lead to a blurry vision (Geurten et al., 2014). Although precise definitions vary slightly, saccades are abrupt turns ranging from 20 to 180 degrees over the course of 31 to 67 ms (Muijres et al., 2015). We defined saccades as turns on the X-Y plane of at least 30 degrees (0.53 radians) in 5 frames (50 ms) or less. We calculated angular velocity using the normalized position vector in the X-Y plane, which projects the X-Y positions on a unit circle. We then calculated the angles between two consecutive position vectors to obtain the angular change in 1 frame (10 ms) for each frame throughout a trajectory. We identified saccades as spikes in angular velocity (Fig. 1D) using a traditional peak finding algorithm from *SciPy*, then selected peaks at least 30 degrees per 50 ms in height.

### Machine Learning Analysis

To prepare our training data we first computed a set of flight patterns: curvature, acceleration, speed at saccade, z-directional speed, distance traveled, and mean speed between saccades. Since the total number of trajectories differed among groups, we sub-sampled a random dataset from the larger group to obtain a dataset with the same data size as the smaller one, as needed. We used ⅔ of our data for training and the rest as test data. We then standardized the data (mean = 0, variance = 1) to reconcile the scaling differences between values.

We compared the data classification accuracy of a linear classifier, Linear Discriminant Analysis (LDA), and a non-linear classifier, the multilayer perceptron (MLP). We constructed a simple two-layer neural network with 16 hidden units and LeakyRelu as the activation function. To optimize the neural network, we used the Adam optimizer at a 0.001 learning rate with gradient descent and other optimization parameters set to default. The neural network was trained using Binary Cross Entropy loss for 50 epochs. We repeated the data subsampling, splitting, training, and testing of classifiers for 100 different seeds. These processes were completed using standard Python (version 3.8.5) packages from Scikit-learn (version 1.1.0), PyTorch (2.3.0+cu121) and NumPy (version 1.24.4).

## RESULTS

### Fruit fly crepuscular activity increases with age and ommatidial diameter under naturalistic lighting

To better understand the effect of ambient brightness and ommatidial diameter on general daily activity, we placed a group of flies in LAMs under an abrupt 12L:12D (Fig. 2A), and another group in a gradual loop that linearly ramps on from 6:00 to 8:00 and off from 18:00 to 20:00 (Fig. 2B). The gradual condition maintained the same circadian period and total luminance while simulating the incremental brightness changes during sunrise and sunset. Assuming flies try to maximize crepuscular activity, we predicted that sensitivity differences due to smaller ommatidia would reduce crepuscular activity primarily in the gradual condition.

Because vinegar flies are crepuscular, most activity occurs during the early and late periods of daylight, highlighted in gray spans of the daily means plots in Fig. 2C, D. As flies aged, their activity increased. Whereas only early activity increased in the abrupt condition, both early and late activity increased significantly in the gradual condition (Fig. 2E), and as a result, showed significantly greater late activity than the abrupt condition starting on day 4 (Fig. 2E).

We then analyzed the relation between ommatidial diameter and mean daily activity (Fig. 2F). Note that we repeated this approach with early and late mean activity as well, but found the same qualitative effects (Fig. S1). We followed a systematic approach to find the best-fitting model predicting mean overall activity as a function of lighting condition, ommatidial diameter, their interaction, and all combinations thereof (Table S1) using OLS. The best-fitting model (adjusted R^2^ = .20, p = .007) with significant predictors represents mean activity as the sum of light condition and the interaction between light condition and ommatidial diameter. Because the light condition is dummy-coded (see methods), this produces a coefficient for the effect of ommatidial diameter on mean activity in each lighting condition. Whereas the coefficients for condition (0.005, p = .208) and the effect of ommatidial diameter in the abrupt condition (4.537, p = .225) were not significant, the effect of ommatidial diameter in the gradual condition (17.013, p = .003) was significant, positive, and over 3 times the effect in the abrupt condition (Fig. 2G). This is consistent with a roughly two-fold increase in Pearson correlations (abrupt: r = .23, p = .286; gradual: r = .48, p = .028).

To understand how the relation between ommatidial diameter and mean activity changes with age, we modeled mean daily activity as the sum of age and the interaction of light condition and ommatidial diameter using a GLMM to account for within subject covariances (predicted means are plotted as dashed lines in Fig. 2F and the results are in Table S1). This model found a significant increase in mean daily activity for with age (0.005 per day, p = 1.13e-19), and corroborated the interactions found in the OLS model after accounting for age-related differences (see slope information at the top of Fig. 2F). Overall, these regression models and bootstrapped mean distributions agree that a gradually changing light schedule—typical of natural crepuscular activity—results in a positive correlation between ommatidial diameter and mean activity, as predicted, and an age-related increase of late activity.

### Fruit flies lower their speed and saccade less frequently during the day if light is scarce

To examine light effects on flight, we next monitored post-eclosion flies behaving in flight cages. We used gradual light changes as above but simulated dim days by programming LEDs to reach a maximum of 25% of the control brightness (Fig. 1F) and kept all other conditions constant between two groups. The light cycle here transitioned gradually from 5:00 to 7:00 (dawn) and from 17:00 to 19:00 (dusk), but the light intensity in the dim cage was always lower. We examined saccades (abrupt turns) during flight, and flight speeds around and between saccades. Overall, the number of saccades per trajectory was unaffected by light level (z= 1.46, p = .145). Most flights had no saccades (Fig. 3B), and the mean angular displacement of saccades stayed comparable between the dim and bright light groups (Fig. 3A, C; z = -1.68, p = .09), except at dusk, where a brighter environment increased angular speed. We also found that in dim light flies saccade more frequently, specifically during the 2 hour period at dawn (Fig. 3F; z = -4.39, p = 1.17e-05).

**Fig. 3.**
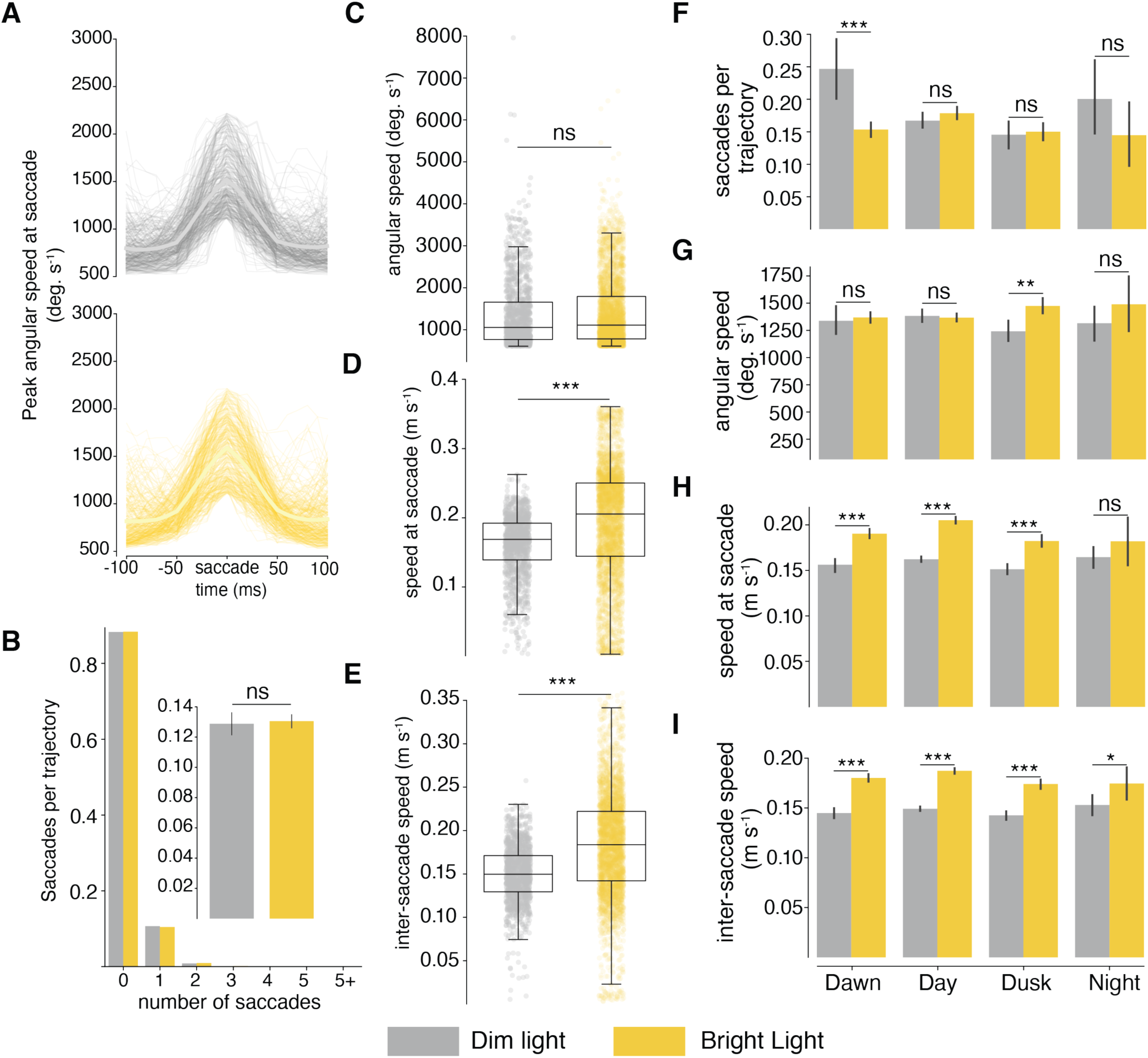
Flight patterns of flies under dim (gray) and bright (yellow) daylight conditions. (A) Random sample of 50 saccades from each group plotted as angular speed in degrees per second 100 ms before and after a saccade. Lighter solid line denotes the mean angular speed. (B) Mean number of saccades per trajectory, (C) mean angular speed at saccade in degrees per second, (D) mean speed within 50 ms around saccade, and (E) translational speed between saccades. For C-E, the data in boxes denotes the 25-75 percentile with the horizontal line representing the median. Whiskers cover q1 ± 1.5* interquartile range. (F-I) Flight patterns under dim (gray) and bright (yellow) daylight conditions at 4 time periods (dawn 5:00-7:00, day 7:00-17:00, dusk 17:00-19:00, night 19:00-5:00). Black error bars represent 95% confidence intervals. Asterisks label significant differences between datasets after running z tests (* for ɑ< 0.05, ** for ɑ <0.01, and *** for ɑ <0.001). A summary of all statistical values can be found on Table S2.

We found robust differences in mean flight speed, where dimmer light made the flies significantly slower both within 50 ms of a saccade (Fig. 3D; z = -14, p = 1.56e-44) and in between saccadic turns (Fig. 3E; z = -19, p = 1.4e-81). In between saccades, the speed difference conditioned by the light environment persisted even after dusk (Fig. 3I, z = -2.4, p = .017), suggesting that prolonged light scarcity also influences behavior at night.

### Smaller flies lower their speed as much as regular flies in dim light

For compound eyes, size directly affects light gathering power, and decreasing light levels might affect smaller individuals more than larger conspecifics. This likely explains why flies with smaller facets were less active during the dim transitions in the LAM experiments. But LAMs largely monitor walking, a less visually demanding behavior than flight. To determine how size affects flight in dim environments, we tested small and large adults in parallel under the same bright light conditions. Here we found the number of saccades per trajectory (Fig. 4B) and the angular speed at saccade (Fig. 4A, C) were slightly higher for the small fly group (z = 2.15, p = .032). For translational speed, large flies were significantly faster during (Fig. 4D; z = -18.03, p = 1.05E-72) and between saccades (Fig. 4E; z = -19.81, p = 2.34E- 87). This trend was consistent — regardless of the time of day, larger flies were faster (Fig. 4G, H, Table S2).

**Fig. 4.**
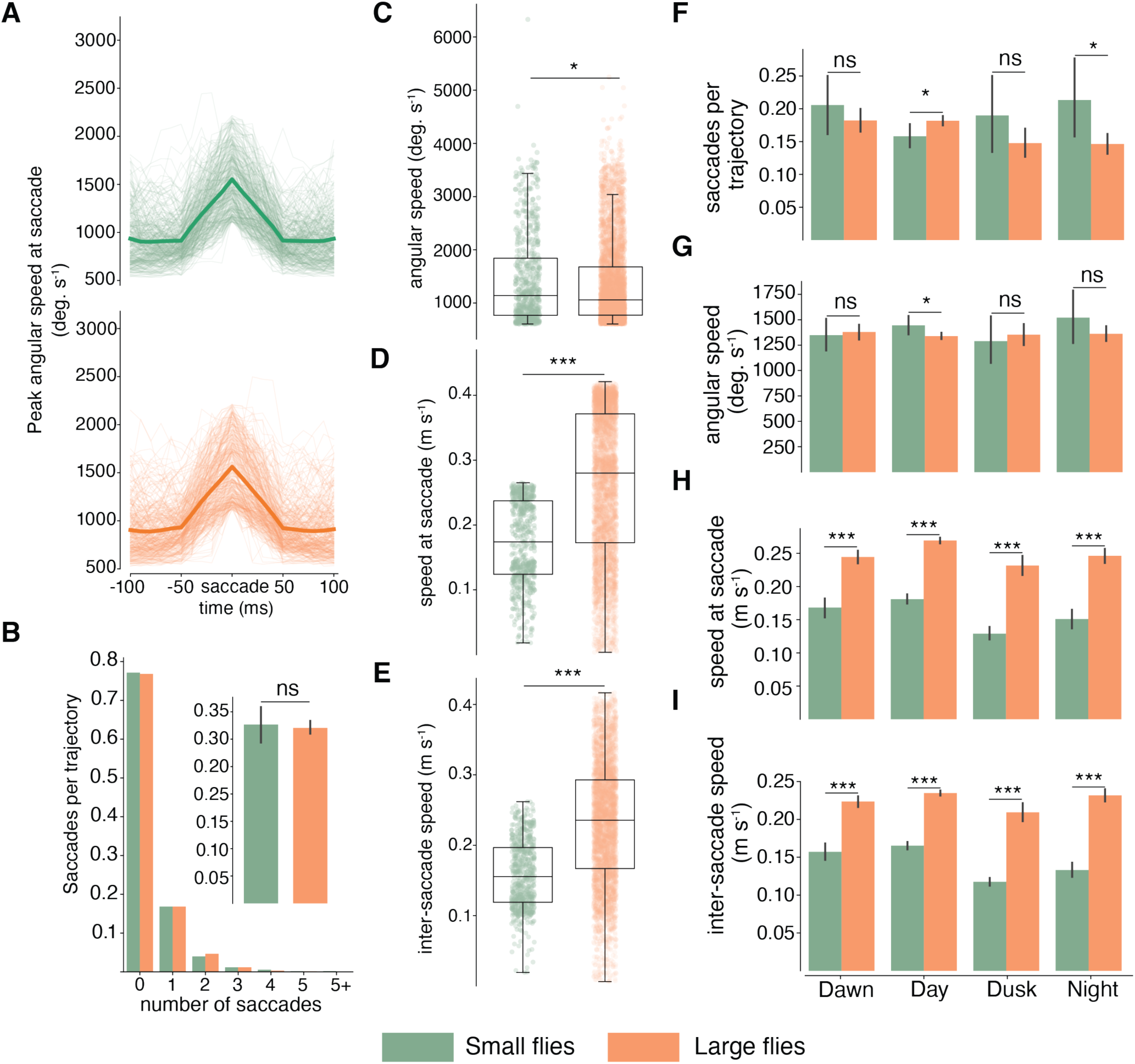
Flight patterns of small (green) and large (orange) flies. (A) Random sample of 50 saccades from each group plotted as angular speed in degrees per second 100 ms before and after a saccade. Lighter solid line denotes the mean angular speed. (B) Mean number of saccades per trajectory, (C) mean angular speed at saccade in degrees per second, (D) mean speed within 50 ms around saccade, and (E) translational speed between saccades. For C-E, the data in boxes denotes the 25-75 percentile with the horizontal line representing the median. Whiskers cover q1 ± 1.5* interquartile range. (F-I) Flight patterns under dim (gray) and bright (yellow) daylight conditions at 4 time periods (dawn 5:00-7:00, day 7:00-17:00, dusk 17:00-19:00, night 19:00-5:00). Black error bars represent 95% confidence intervals. Asterisks label significant differences between datasets after running z tests (* for ɑ< 0.05, ** for ɑ <0.01, and *** for ɑ <0.001). A summary of all statistical values can be found on Table S2.

Total saccade rate did not vary significantly with fly size (Fig. 3B; z = -0.38, p = .705), unless we considered night flights alone. We identified only 253 night trajectories for small flies, but 3150 for large flies, yet smaller individuals saccade more frequently at night (Fig. 3F; z = 2.40, p = 0.016).

### Small flies can compensate for their limited optics if more light is available

Lower flight speeds in small individuals might be due to either reduced light capture, or diminished wing or body size restricting flight abilities. We wondered if increasing the light levels could compensate for having smaller optics. To test this we illuminated the experimental cage with a second set of LEDs (a treatment we call Bright 2X) and tested both fly sizes in parallel. Under these conditions, where the insects were experiencing roughly twice the total brightness they did in previous experiments, small flies sped up significantly around saccades (Fig. 5C; z = -14.14, p = 2.29E-45) and between them (Fig. 5D; z = -15.94, p = 3.62E-57) compared to previous experiments of small flies in regular bright light. The mean speed changed roughly by a 1.3-fold, from 0.172 to 0.225 m/s for speed around saccades, and from 0.157 to 0.205 for mean speed between saccades, a difference we expect to be of biological relevance.

**Fig. 5.**
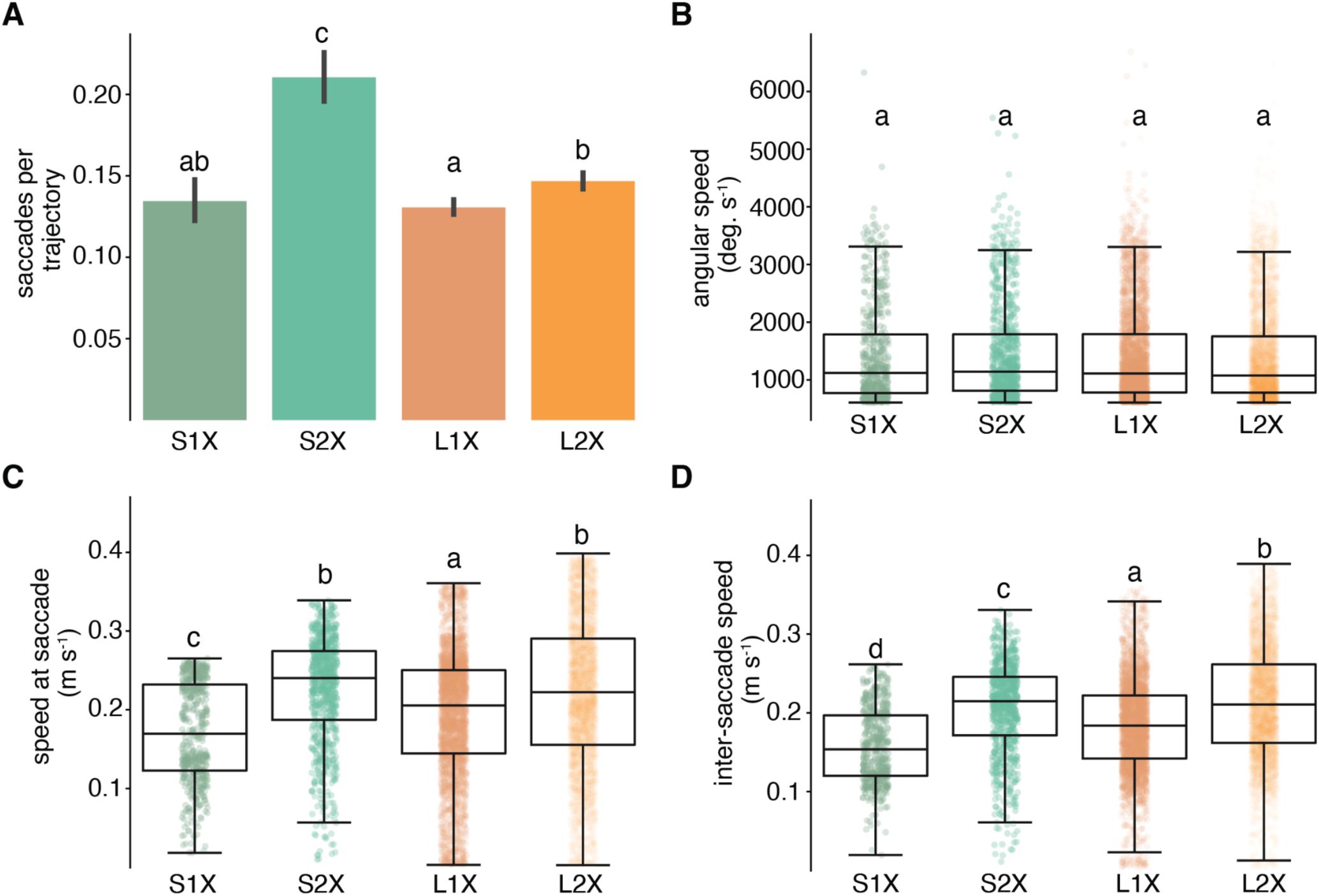
Small and large flies in regular vs brighter light environments. Number of saccades per trajectory (A), angular speed in degrees per second (B), speed within 50 ms around saccades (C) and speed in between saccades (D) were compared among four treatment groups: small flies in bright light (green), small flies in brighter (2X) light (bright green), large flies in bright light (orange), and large flies in brighter (2X) light (bright orange). The vertical lines in the bar plots (A) mark the 95% confidence interval. Data in boxes (B-D) includes the 25-75 percentile with a horizontal line denoting the median. Whiskers cover q1 ± 1.5* interquartile range. Letters on top of each dataset label significant differences between groups after running z tests. A summary of all statistical values can be found on Table S2.

Remarkably, not only were small flies faster when more light became available (S2X flies), but they were faster (mean= 0.225 m/s) than large flies under regular brightness (L1X, mean= 0.194) — that is, those that only had one set of lights during the day (Fig. 5C; z = -10.09, p = 5.92E-24 and Fig. 5D; z = -10.13, p = 4.02E-24). These results suggest light capture was the factor limiting flight speed for smaller individuals.

An increase in daylight also led to a slight increase in the number of saccades per trajectory, especially for small flies, for which the percentage of flight containing at least one saccade went from 14 to 23% (Fig. 5A; z = -8.32, p = 8.52E-17), while the angular speed was unaffected (Fig. 5B, z = - 0.18, p = .859, Table S3).

### Translational but not angular speed is needed to classify fly treatment groups

After finding experimental speed differences between groups of flies, we investigated the existence of non-linear interactions between features of flight trajectories using machine learning as described in above. Because of the complex relationship between body size, ambient light, and flight performance, we did not expect these variables to influence flight in a direct (linear) manner. We examined the performance of LDA vs MLP classifiers in separating fly groups by age, size, and light niche using flight trajectory features like speed, distance, and acceleration.

As predicted, MLP consistently outperformed the LDA in separating light, size, and age (Fig. 6; 8- 18% increased accuracy), suggesting that the MLP extracts useful non-linear feature differences to improve data classification accuracy. We then conducted restriction analyses on the LDA and MLP, where we dropped flight dynamics features and studied how the classification results degraded. For both classifiers, their performance decreased the most when three features- inter-saccade translational speed, translational speed at saccade, and distance traveled during saccade-were excluded from the analysis. This analysis suggests that these three features are fundamental indicators of the differences in flight dynamics that result from modifying a treatment variable like light, body size and fly age. Notably, the specific non-linear relationship between each condition (age, size, light niche) and flight behavior remains unknown, but our results indicate that it is primarily the translational rather than rotational aspects of flight that vary with age, size, and light level.

**Fig. 6.**
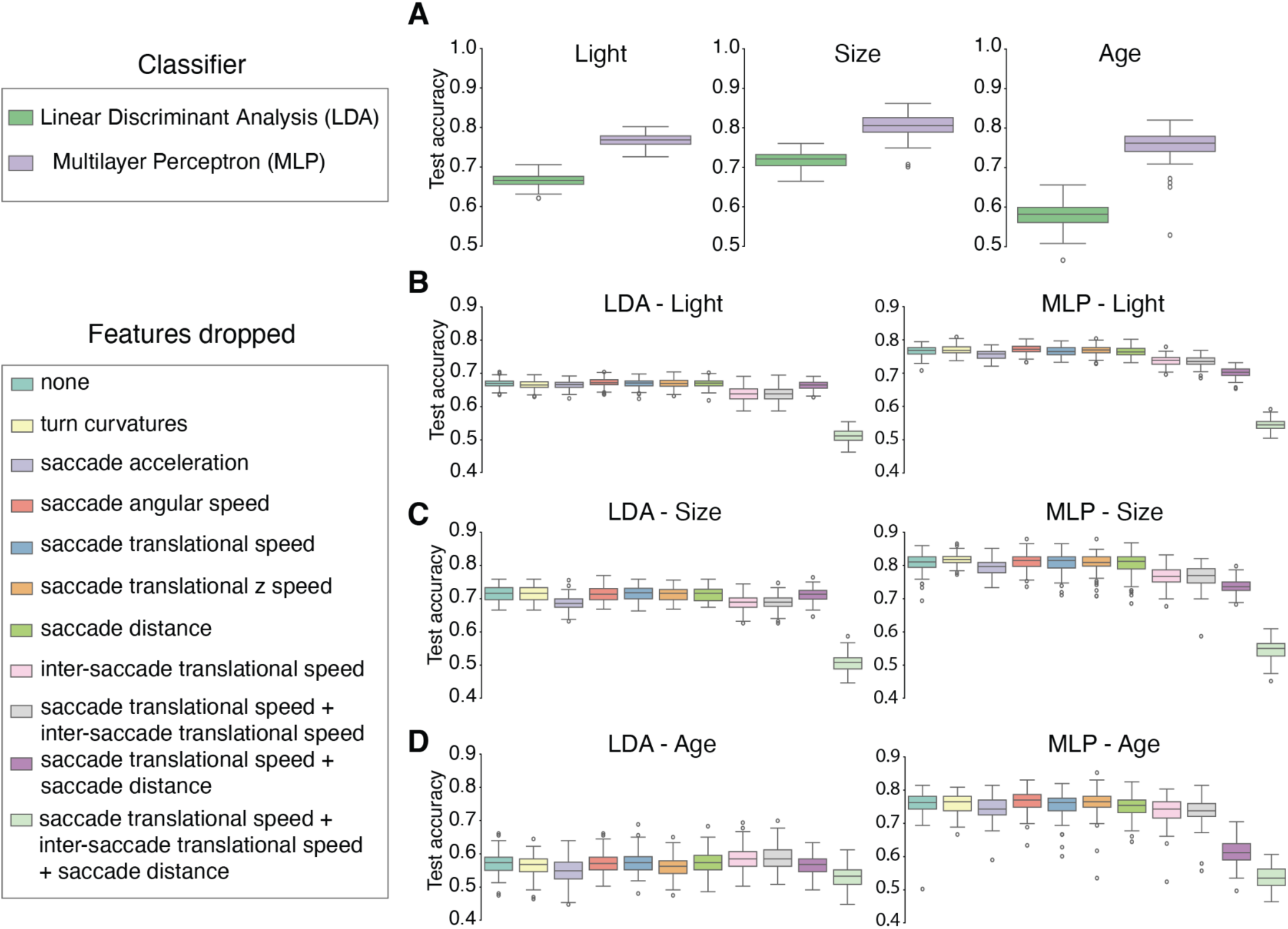
Machine learning analysis comparing two methods to separate fly groups based on their saccades. (A) Classification accuracy of a linear classifier (LDA) compared to a non-linear classifier (MLP) for flies in bright vs. dim light (left), small vs large flies (middle) and day 1 vs day 4 flies (right) data groups. (B-D) Ablation study testing flight feature importance for accurate treatment classification. Classification accuracy based on each condition light (C), size (D), and age (D) groups decreased by a different magnitude depending on the specific feature(s) removed from the network.

## DISCUSSION

Low photon catch threatens reliable vision and the behaviors that depend on it. This becomes an issue not just in dim environments, but when scenes move quickly, or optical elements are small. To improve catch and image quality, animals, like photographers, must either devote resources to larger optics, sacrifice some form of acuity, or strategically alter their behaviors. Like many other insects, fruit flies operate with small eyes, necessitated by their small size, but a strong need for dependable images to control safe and effective flight. We examined activity and flight behavior to determine if vinegar flies alter their activity or flight patterns to compensate for low photon catch.

During flight, fruit flies may saccade more or less frequently depending on the visual scene. For instance, compared to panoramic views, which mostly elicit smooth turns, flies saccade consistently when tracking a moving vertical bar (Fox & Frye, 2013; Mongeau & Frye, 2017), described also in walking insects (Geurten et al., 2014). In a naturalistic scene, flies’ saccadic behavior is modulated by the dynamics and speed of moving objects (Mongeau & Frye, 2017). But in the absence of distractive visual cues, we were able to study how light and body size alone influence flight dynamics. The differences we found in saccadic turns between treatment groups were not as strong as those in smooth flight speed. Notably, only the translational speed and not the angular change during a saccade was modulated by size and light levels. We confirmed this trend using neural network training, where we achieved optimal test accuracy to classify data only when we included the translational speed features (Fig. 6).

Within the same insect species, body size does not dictate speed in a straightforward manner. While larger individuals may have higher muscular mass, the efficiency of these muscles and other factors, such as metabolic rate and wing loading, play crucial roles in flight performance (Marden, 2000). Smaller individuals often exhibit high specific metabolic rates, allowing for quick bursts of energy and agile movements. Additionally, ecological adaptations within a species may result in variations, where some individuals may specialize in sustained flight, while others excel in quick, precise movements for specific tasks (Dudley, 2002). Even slight variations in eye size among closely related Drosophilids can have dramatic consequences for their vision (Buffry et al., 2024). The present study shows how a poor larval diet may alter the flight dynamics of adult flies, with possible consequences, such as predation risk or reduced mating success.

Although at the optical level small fruit flies sacrifice their ability to capture light by having fewer and smaller ommatidia, they recover this ability neurally by sacrificing temporal acuity (Currea et al., 2018). Here, we demonstrated that smaller ommatidia result in less general activity under a gradually changing light schedule (Fig. 2). And using a free flight assay, we found that small flies fly more slowly than larger ones at every time interval analyzed (dawn, day, dusk, night), but maintained generally equal saccade magnitudes (Fig. 4). Moreover, all flies flew faster in brighter light, with small flies comparable to large flies in a brighter context (Fig. 3). In combination, these results imply that temporal summation used to recover contrast sensitivity has higher-order consequences on behavior and ecology. In dim conditions fewer photons reach the retina, and without neural tricks this results in poor contrast sensitivity. Photoreceptors generally longer to respond to visual stimuli, and often pool information spatially to improve this sensitivity. Slower response times impact the ability to react quickly, such as avoiding obstacles or predators, and reduced spatial acuity sacrifices object recognition, edge detection, or identification of other animals. We found dim daylights slowed flies, and the trend persisted even after sunset (Fig. 3I). This suggests that flies housed under limited light acclimate their flight speed and are unlikely to take on faster flights. Effectively, limiting light made flies behave similar to small flies, suggesting a strong connection between visual optics and environmental luminosity in flight strategies. Lab reared flies are traditionally well-fed and raised in rooms with standard but abruptly transitioning light cycles. Part of what makes *Drosophila* such a useful model organism is the ease and reproducibility of its care. But understanding natural behaviors often requires replicating important environmental conditions in the lab. Generating more naturalistic light cycles indoors is now straightforward and inexpensive. Further, typical lab diets generate large individuals with little of the size variation found in the wild, but this is easily replicated in lab populations. As we show here, both lighting and size affect activity and flight behavior, and for some research questions, these are crucial for generalizing laboratory findings to natural populations.

## Acknowledgments

We thank members of the Theobald Laboratory at FIU, and the Tarokh Laboratory at DU. This project was funded by the FLAP (Fast Lexicographic Agile Perception) MURI (Multidisciplinary Research Initiative) grant by the Department of Defense. We also thank Dr. Mark Frye and our collaborators from the FLAP MURI group for their feedback on our current study. This content is solely the responsibility of the authors and does not represent the views of the DoD.

## Competing Interests

The authors declare no competing interests.

## Authors contributions

EB conceptualization, methodology, data gathering, writing, visualization, data curation, project administration. HY conceptualization, data analysis, writing, visualization. PC conceptualization, methodology, data gathering, writing, visualization, data curation. YS methodology, review and editing. RP methodology, review and editing. SS methodology, review and editing, project administration, funding acquisition. VT conceptualization, data analysis, writing, project administration, funding acquisition. JT conceptualization, methodology, writing, project administration, funding acquisition.

## Funding

Financial support was provided by the National Science Foundation (NSF IOS-1750833 to JCT), and U.S. Air Force Office of Scientific Research (AFOSR MURI awarded FA9550-22-1-0315 to JCT, VT, and SS).

## Data availability

Data and code are available at https://doi.org/10.6084/m9.figshare.26122231.v2 and 10.6084/m9.figshare.26210627

## Supplementary Resources

**Figure S1.**
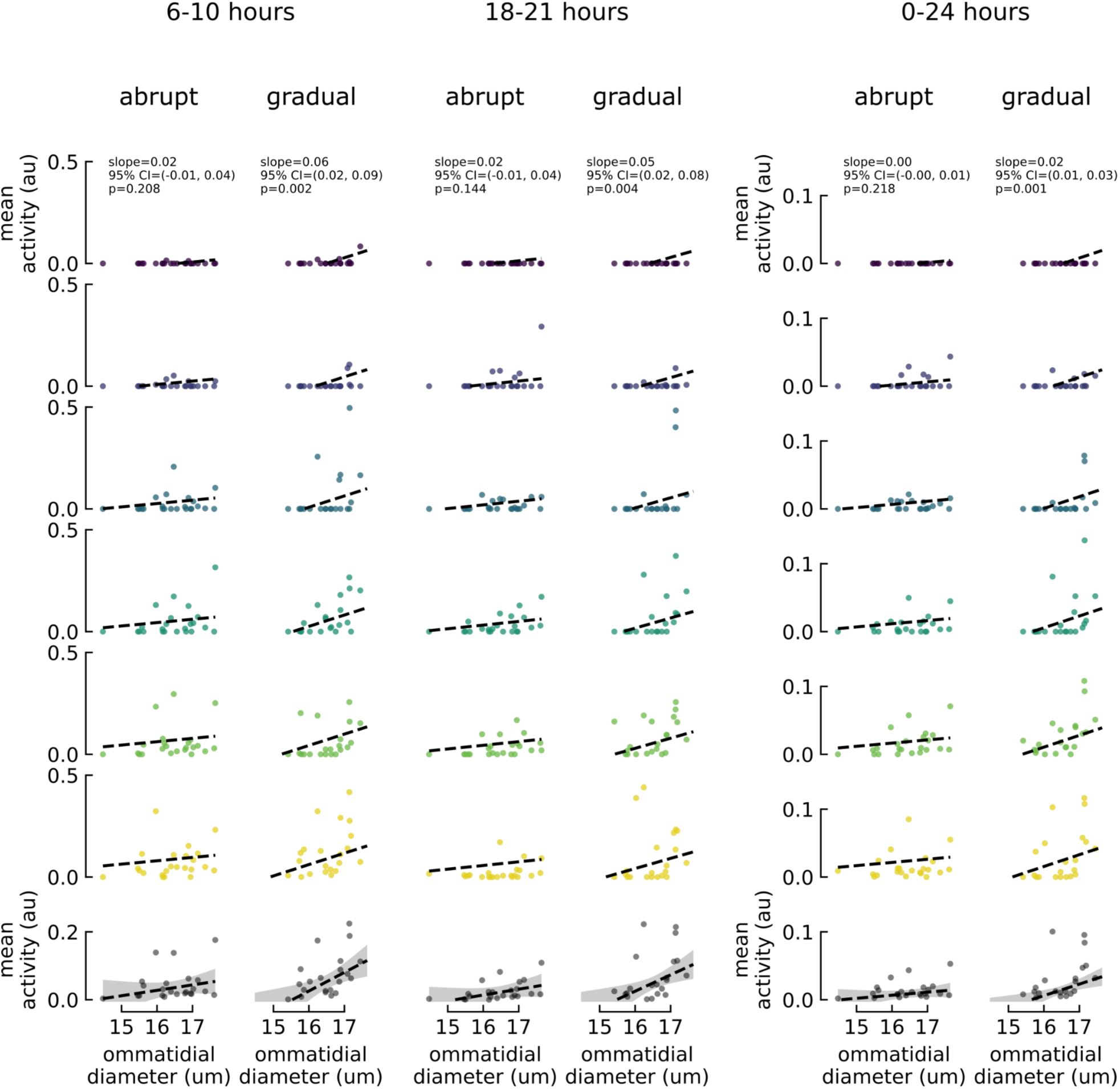
Mean daily activity and ommatidial diameter during 3 different time spans: 6-10 (early), 18-21 (late), and 0-24 (all day) hours. Mean daily activity is normalized to the maximum bin-averaged activity (1.69 events per minute) and therefore expressed in arbitrary units (au). The early and late columns share the same y-axis. Age increases for each row from top to bottom with mean activity across all days plotted in the bottom row. The slope parameter, its 95% CI of the mean, and p-value produced by the GLMM are printed at the top of each column. Predicted means are plotted as dashed lines and the gray error bands in the bottom row represent the predicted 95% CI of the mean of an OLS model regressing mean activity on condition and the interaction of condition and ommatidial diameter. Note that all three spans and the resulting regressions produce nearly equivalent qualitative results.

**S2.**
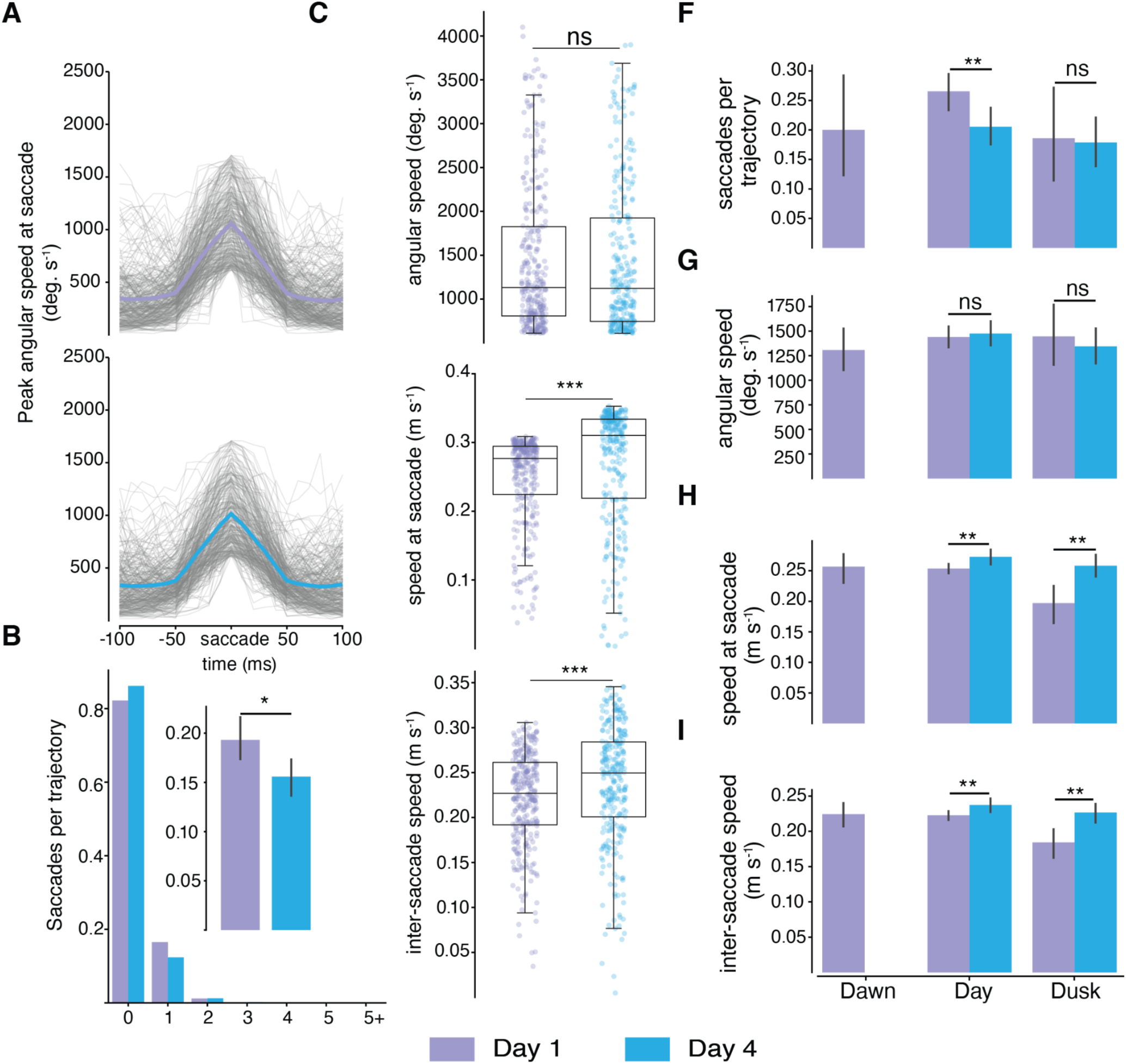
Flight patterns of day 1 (violet) and day 4 (blue) flies. (A) Sample of 50 saccades from each group plotted as angular speed in degrees per second 100 ms before and after a saccade. Lighter solid line denotes the mean angular speed. (B) Mean number of saccades per trajectory, (C) mean angular speed at saccade in degrees per second, (D) mean speed within 50 ms around saccade, and (E) translational speed between saccades. For C-E, the data in boxes denotes the 25-75 percentile with the horizontal line representing the median. Whiskers cover q1 ± 1.5* interquartile range. (F-I) Flight patterns for day 1 (violet) and day 4 (blue) flies at 3 time periods (dawn 5:00-7:00, day 7:00-17:00, dusk 17:00-19:00). Night 19:00-5:00 was omitted since no trajectories were identified at this time range. Black error bars represent 95% confidence intervals. Asterisks label significant differences between datasets after running Z tests.

**Table S1.**
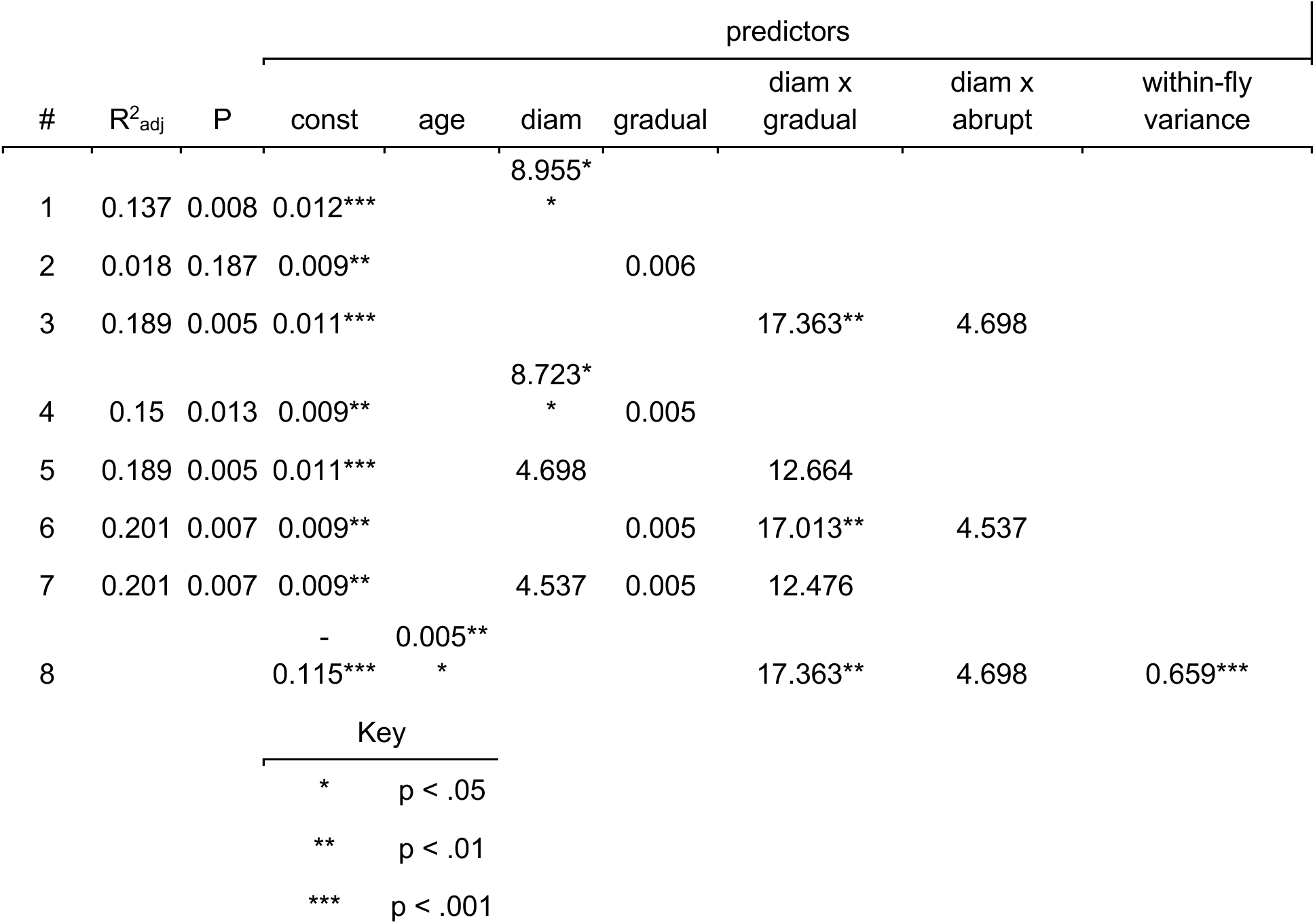
Regression models of overall and daily mean activity. Results of OLS regression models with different combinations of the predictors. Each row is a different equation model providing the proportion of explained variance adjusted for the number of predictors involved (R^2^_adj_), the significance of the model (P), and the resultant coefficient for each predictor involved and its corresponding significance represented by asterisks according to the key. Empty predictor cells mean that the predictor was excluded from that model. The inclusion of both diam and diam x gradual predictors results in loss of significance for either, suggesting multicollinearity between the two predictors. In other words, the significant effect of diameter in models 1 and 4 is primarily due to the significant effect of diameter in the gradual condition. This is replicated each time both terms are included as predictors (models 5 and 7), so we avoided models with both terms in our analysis.

**Table S2.** Statistical analysis results related to Figures 2,3, and 4. Top row denotes the time period where treatment groups were analyzed, where ‘Total Daily’ refers to the entire day. Orange rows indicate the treatment groups compared and gray rows refer to the parameter for which a z test was conducted.

|  | Total Daily |  | Dawn |  | Day |  | Dusk |  | Night |  |
| --- | --- | --- | --- | --- | --- | --- | --- | --- | --- | --- |
| Dim vs Bright |  |  |  |  |  |  |  |  |  |  |
| number of trajectories |  |  |  |  |  |  |  |  |  |  |
| total | 7433 | 21508 | 458 | 5806 | 5371 | 9445 | 1285 | 4327 | 319 | 1930 |
| saccade translational speed (m/s) |  |  |  |  |  |  |  |  |  |  |
| mean | 0.160 | 0.194 | 0.156 | 0.190 | 0.162 | 0.205 | 0.151 | 0.12 | 0.164 | 0.155 |
| z score | -14.00 |  | -4.38 |  | -15.56 |  | -4.95 |  | 0.81 |  |
| p value | 1.56E-44 |  | 1.17E-05 |  | 1.34E-54 |  | 7.42E-07 |  | 0.4176 |  |
| saccade angular speed (degrees / s) |  |  |  |  |  |  |  |  |  |  |
| mean | 1353.8 | 1398.2 | 13377 | 1368.3 | 1382.1 | 1366.6 | 1240.9 | 1475 | 1315.1 | 1524.2 |
| z score | -1.68 |  | -0.41 |  | 0.47 |  | -3.28 |  | -1.76 |  |
| p value | 0.094 |  | 0.681 |  | 0.640 |  | 0.001 |  | 0.078 |  |
| between saccade speed |  |  |  |  |  |  |  |  |  |  |
| mean | 0.148 | 0.181 | 0.145 | 0.180 | 0.149 | 0.187 | 0.143 | 0.14 | 0.153 | 0.165 |
| z score | -19.13 |  | -6.32 |  | -18.27 |  | -6.87 |  | -1.50 |  |
| p value | 1.40E-81 |  | 2.67E-10 |  | 1.42E-74 |  | 6.63E-12 |  | 0.1326 |  |
| number of saccades per trajectory |  |  |  |  |  |  |  |  |  |  |
| mean | 0.17 | 0.16 | 0.25 | 0.15 | 0.17 | 0.18 | 0.15 | 0.15 | 0.20 | 0.12 |
| z score | 1.46 |  | 4.42 |  | -1.48 |  | -0.35 |  | 3.01 |  |
| p value | 0.1446 |  | 9.69E-06 |  | 0.1399 |  | 0.7229 |  | 0.0026 |  |
| Small vs Large |  |  |  |  |  |  |  |  |  |  |
| number of trajectories |  |  |  |  |  |  |  |  |  |  |
| total | 3123 | 21289 | 384 | 2603 | 2270 | 13904 | 216 | 1632 | 253 | 3150 |
| saccade translational speed (m/s) |  |  |  |  |  |  |  |  |  |  |
| mean | 0.172 | 0.261 | 0.168 | 0.244 | 0.181 | 0.269 | 0.129 | 0.22 | 0.151 | 0.246 |
| z score | -18.03 |  | -6.25 |  | -14.80 |  | -5.78 |  | -6.09 |  |
| p value | 1.05E-72 |  | 4.15E-10 |  | 1.57E-49 |  | 7.65E-09 |  | 1.14E-09 |  |
| saccade angular speed (degrees / s) |  |  |  |  |  |  |  |  |  |  |
|  | 1425.9 | 1347.7 | 1348.8 | 1379.1 | 1444.3 | 1338.7 | 1288.6 | 135.1 | 1520.5 | 1361.7 |
| z score | 2.15 |  | -0.32 |  | 2.40 |  | -0.47 |  | 1.36 |  |
| p value | 0.032 |  | 0.751 |  | 0.016 |  | 0.637 |  | 0.174 |  |
| between saccade speed |  |  |  |  |  |  |  |  |  |  |
| mean | 0.157 | 0.231 | 0.157 | 0.224 | 0.165 | 0.235 | 0.117 | 0.209 | 0.133 | 0.232 |
| z score | -19.81 |  | -7.23 |  | -15.41 |  | -6.51 |  | -7.96 |  |
| p value | 2.34E-87 |  | 4.88E-13 |  | 1.45E-53 |  | 7.30E-11 |  | 1.75E-15 |  |
| number of saccades per trajectory |  |  |  |  |  |  |  |  |  |  |
| mean | 0.171 | 0.174 | 0.206 | 0.182 | 0.158 | 0.182 | 0.190 | 0.148 | 0.213 | 0.146 |
| z score | -0.38 |  | 0.92 |  | -2.31 |  | 1.33 |  | 2.40 |  |
| p value | 0.705 |  | 0.360 |  | 0.021 |  | 0.184 |  | 0.016 |  |
| Day1 vs Day4 |  |  |  |  |  |  |  |  |  |  |
| number of trajectories |  |  |  |  |  |  |  |  |  |  |
| total | 1216 | 1399 | 130 | 0 | 946 | 901 | 140 | 498 | 0 | 0 |
| saccade translational speed (m/s) |  |  |  |  |  |  |  |  |  |  |
| mean | 0.249 | 0.268 | 0.256 | n/a | 0.254 | 0.273 | 0.197 | 0.258 | n/a | n/a |
| z score | -2.99 |  | n/a |  | -2.70 |  | -3.21 |  | n/a |  |
| p value | 0.003 |  | n/a |  | 6.92E-03 |  | 1.32E-03 |  | n/a |  |
| saccade angular speed (degrees / s) |  |  |  |  |  |  |  |  |  |  |
| mean | 1430.1 | 1433.3 | 1308.6 | n/a | 1441.0 | 1475.7 | 1446.3 | 1345.3 | n/a | n/a |
| z score | -0.05 |  | n/a |  | -0.41 |  | 0.54 |  | n/a |  |
| p value | 0.963 |  | n/a |  | 0.679 |  | 0.590 |  | n/a |  |
| between saccade speed |  |  |  |  |  |  |  |  |  |  |
| mean | 0.220 | 0.234 | 0.224 | n/a | 0.223 | 0.237 | 0.184 | 0.227 | n/a | n/a |
| z score | -2.91 |  | n/a |  | -2.61 |  | -3.00 |  | n/a |  |
| p value | 0.004 |  | n/a |  | 8.93E-03 |  | 2.71E-03 |  | n/a |  |
| number of saccades per trajectory |  |  |  |  |  |  |  |  |  |  |
| mean | 0.249 | 0.196 | 0.200 | n/a | 0.265 | 0.205 | 0.186 | 0.179 | n/a | n/a |
| z score | 2.79 |  | n/a |  | 2.60 |  | 0.16 |  | n/a |  |
| p value | 0.005 |  | n/a |  | 0.009 |  | 0.875 |  | n/a |  |

**Table S3.** Table S2. Statistical analysis results related to Figure 5. For simplicity, we labeled the treatment groups as follows: L1X = large flies in bright light, L2X = large flies in 2X bright light, S1X = small flies in bright light, S2X = large flies in 2X bright light. Orange rows indicate the treatment groups compared and gray rows refer to the parameter for which a z test was conducted.

|  | L1X vs S2X |  | L1X vs L2X |  | S1X vs S2X |  | L2X vs S2X |  |
| --- | --- | --- | --- | --- | --- | --- | --- | --- |
|  | saccade translational speed (m/s) |  |  |  |  |  |  |  |
| mean | 0.194 | 0.225 | 0.194 | 0.220 | 0.172 | 0.225 | 0.220 | 0.225 |
| z score | -10.09 |  | -12.85 |  | -14.14 |  | -1.33 |  |
| p value | 5.92E-24 |  | 8.69E-38 |  | 2.29E-45 |  | 0.183 |  |
|  | saccade angular speed (degrees/s) |  |  |  |  |  |  |  |
| mean | 1398.2 | 1434.3 | 1398.2 | 1394.9 | 1425.9 | 1434.3 | 1394.9 | 1434.3 |
| z score | -1.14 |  | 0.17 |  | -0.18 |  | -1.24 |  |
| p value | 0.2540 |  | 0.8627 |  | 0.8586 |  | 0.216 |  |
|  | between saccade speed |  |  |  |  |  |  |  |
| mean | 0.181 | 0.205 | 0.181 | 0.210 | 0.157 | 0.205 | 0.210 | 0.205 |
| z score | -10.13 |  | -18.72 |  | -15.94 |  | 1.93 |  |
| p value | 4.02E-24 |  | 3.71E-78 |  | 3.62E-57 |  | 0.053 |  |
|  | number of saccades per trajectory |  |  |  |  |  |  |  |
| mean | 0.16 | 0.28 | 0.16 | 0.18 | 0.17 | 0.28 | 0.18 | 0.28 |
| z score | -13.14 |  | -5.26 |  | -8.32 |  | -10.02 |  |
| p value | 2.02E-39 |  | 1.42E-07 |  | 8.52E-17 |  | 1.20E-23 |  |

## Notes

### Competing Interest Statement

The authors have declared no competing interest.

